# Neural dysfunction correlates with heat coma and CT_max_ in *Drosophila* but does not set the boundaries for heat stress survival

**DOI:** 10.1101/844316

**Authors:** Lisa B. Jørgensen, R. Meldrum Robertson, Johannes Overgaard

## Abstract

When heated, insects loose coordinated movement followed by the onset of heat coma (CT_max_). These phenotypes are popular measures to quantify inter- and intraspecific differences in insect heat tolerance, and CT_max_ correlate well with current species distributions. Here we examined the function of the central nervous system (CNS) in five species of *Drosophila* with different heat tolerances, while they were exposed to either constant high temperature or a gradual increasing temperature (ramp). Tolerant species were able to preserve CNS function at higher temperatures and for longer durations than sensitive species and similar differences were found for the behavioral indices (loss of coordination and onset of heat coma). Furthermore, the timing and temperature (constant and ramp exposure, respectively) for loss of coordination or complete coma coincided with the occurrence of spreading depolarisation (SD) events in the CNS. These SD events disrupt neurological function and silence the CNS suggesting that CNS failure is the primary cause of impaired coordination and heat coma. Heat mortality occurs soon after heat coma in insects and to examine if CNS failure could also be the proximal cause of heat death, we used selective heating of the head (CNS) and abdomen (visceral tissues). When comparing the temperature causing 50% mortality (LT_50_) of each body part to that of the whole animal, we found that the head was not particularly heat sensitive compared to the abdomen. Accordingly, it is unlikely that nervous failure is the principal/proximate cause of heat mortality in *Drosophila*.

**Summary statement:** Hyperthermic failure of the *Drosophila* central nervous system causes heat coma, a phenotype varying in temperature between drosophilids, but neural failure is likely not the primary cause of heat mortality.

## Introduction

Thermal tolerance is arguably among the most important traits in defining the biogeographical distribution of ectothermic species (Addo-Bediako *et al*., 2000; Sunday *et al*., 2014). This is also the case for insects (Gaston & Chown, 1999; Vorhees *et al*., 2013), including *Drosophila* where tolerance to both low and high temperature shows a high correlation to the current species distributions (Andersen *et al*., 2015; Jørgensen *et al*., 2019; Kellermann *et al*., 2012; Kimura, 2004). In the case of insect cold tolerance there is a general understanding of the processes causing cold coma and cold mortality (Andersen *et al*., 2018; Bayley *et al*., 2018; Koštál *et al*., 2004; MacMillan & Sinclair, 2011), and many physiological adaptations that underlie differences in cold tolerance between species and populations have been uncovered (Feder & Hofmann, 1999; Overgaard & MacMillan, 2017; Sinclair *et al*., 2003; Yi & Lee, 2004; Zachariassen, 1985). In contrast, it is generally less clear which physiological perturbations cause heat coma and heat mortality, and accordingly there is a poorer understanding of the adaptations that result in intra- and interspecific variations in insect heat tolerance (but see Bowler (2018) and Neven (2000)).

Heat tolerance of insects and other ectotherms is typically measured by recording the onset of characteristic behaviours (or endpoints) during heat exposure. These measures include the loss of equilibrium or righting response, onset of spasms, entry into a comatose state or heat mortality (Cowles & Bogert, 1944; Lutterschmidt & Hutchison, 1997a; Lutterschmidt & Hutchison, 1997b; Terblanche *et al*., 2011). The term ‘CT_max_’ (critical thermal maximum) is frequently and indiscriminately used for all of these endpoints although the different behavioural phenotypes represent the responses to different intensities or durations of heat stress. Thus, mortality is most often preceded by a progressive loss of motor-control (Friedlander *et al*., 1976; Gladwell *et al*., 1975; Lutterschmidt & Hutchison, 1997a) and some of the endpoints, such as heat coma, can be reversed if the animal is removed from the heat stress immediately after the endpoint is observed (Fraenkel, 1960; Hamby, 1975; Heath *et al*., 1971; Martinet *et al*., 2015; Rodgers *et al*., 2010, but see O’Sullivan *et al*., (2017)). It can be difficult to discriminate the heat coma and heat death (Larsen, 1943; Mellanby, 1954), as the rate of heat injury accumulation responds strongly to small changes in temperature. Accordingly, slightly longer exposures to high temperatures than those causing coma can result in the accumulation of lethal amounts of heat injury (Bigelow, 1921; Jørgensen *et al*., 2019; Kingsolver & Umbanhowar, 2018).

There are a number of physiological dysfunctions that have been suggested to cause heat coma and heat mortality in insects. These include a mismatch between demand and supply of oxygen to active tissues (described in the hypothesis of oxygen and capacity limited thermal tolerance – OCLTT) (Pörtner, 2001), hemolymph hyperkalaemia which would impair muscle function (Gladwell, 1975; Gladwell *et al*., 1975; O’Sullivan *et al*., 2017), cellular heat injury to the membranes (Bowler, 1981; Bowler, 2018; Bowler *et al*., 1973; Hazel, 1995) and breakdown of central nervous function (Hamby, 1975; Larsen, 1943; Prosser & Nelson, 1981; Robertson, 2004). The evidence to support acute heat failure or mortality due to oxygen limitations is not strong for terrestrial insects (Klok, 2004; Mölich *et al*., 2013; Verberk *et al*., 2015) and there is also limited support for hemolymph hyperkalaemia as the proximal cause of heat coma/mortality (O’Sullivan *et al*., 2017). Accordingly, the strongest candidate mechanisms underlying heat coma are tied to breakdown of nervous function. Silencing of nervous function has been observed in heat exposed fruit flies and locusts where heat stress causes a spreading depolarisation (SD) in the central nervous system (CNS) (Money *et al*., 2009; Robertson, 2004; Rodgers *et al*., 2007). Spreading depolarisation is triggered by failure to maintain ion gradients between the intra- and extracellular compartments within the CNS, which results in depolarization of neurons and glial cells and a surge of potassium ions in the extracellular space of the brain, preventing neural activity (Robertson, 2004; Robertson et al. (submitted); Spong et al., 2016). Furthermore, studies have shown that inter- and intraspecific differences in cold coma are highly correlated with the loss of CNS function in insects (Andersen et al., 2018; Robertson et al., 2017). Given the similarity in the behavioural phenotypes of heat and cold coma there is an obvious possibility that the onset of heat coma is also caused by CNS failure in insects.

In most insects, heat mortality follows closely after the onset of heat coma (Mellanby, 1954) and the hypothesis about hyperthermic loss of CNS function could therefore also be extended to be the proximal cause of heat mortality. In goldfish, heating either the cerebellum or the water caused similar behavioural responses, that progressed from hyperactivity to coma (Friedlander *et al*., 1976). A recent study revisited the work of Friedlander *et al*., and here the authors selectively cooled the brain of Atlantic cod while the fish were subjected to heat stress, and found that this resulted in increased heat tolerance (measured as loss of equilibrium), compared to controls and instrumented controls (Jutfelt *et al*., 2019). Accordingly, it appears that controlling the temperature of the CNS can mimic whole-animal exposure to a specific temperature.

In the present study we used a comparative study system of five *Drosophila* species with pronounced interspecific differences in heat tolerance. The most heat sensitive species goes into coma at a temperature 6°C lower than the most tolerant species in a ramping assay, and similarly the constant temperature estimated to cause onset of coma after a 1-hour exposure is almost 6°C lower in the sensitive species compared to the most heat tolerant species used here (Jørgensen *et al*., 2019). To investigate the relation between neural dysfunction and the two behavioural heat stress phenotypes, loss of coordinated movement (T/t_back_) and onset of heat coma (T/t_coma_), we measured DC potentials in the central nervous system of the five species during heat exposure to record spreading depolarisation as an indication of neuronal failure. These experiments were performed with both gradual heating (a dynamic ramping assay) and constant (static) heat exposure to constant temperature. The loss of coordinated movement, the onset of heat coma and heat mortality occur in rapid succession in many insects. To examine if the onset of heat mortality is caused proximately by failure in the CNS, we designed a simple experiment in which we compare the heat sensitivity of flies that are heated over their entire body with specimens heated specifically in the head (CNS) or abdomen (visceral tissues). This experiment was performed in three of the *Drosophila* species and was designed to evaluate if some body sections (head with primarily neuronal tissue vs abdomen with primarily visceral tissue) were more sensitive to heat stress than others.

## Materials and methods

### Experimental animals

Five species of *Drosophila* (*D. immigrans*, Sturtevant 1921; *D. subobscura*, Collin 1936; *D. mercatorum*, Patterson and Wheeler 1942; *D. melanogaster*, Meigen 1830 and *D. mojavensis*, Patterson 1940) were used in this study. The least heat tolerant species *D. immigrans* can survive 35.4°C for 1 hour while the most tolerant species *D. mojavensis* can survive 41.2°C for 1 hour (Jørgensen *et al*., 2019) and collectively these five species represent a broad range of heat tolerances within *Drosophila*. Flies were reared and maintained under common garden conditions in 250-mL bottles containing 70 mL of oat-based Leeds medium (see Andersen et al. (2015)) in a 19°C room with constant light. Maintenance bottles with adults that parented the experimental flies were changed twice a week, and newly eclosed adults from rearing bottles were collected and transferred to fresh vials with fly medium every 1-3 days. Experimental flies were produced by transferring a tablespoon of used medium (including eggs) to another 250-mL bottle with 70 mL new medium. 2-4 days post-eclosion flies were anaesthetised with CO_2_, sexed and female flies were moved to new medium vials, and allowed to recover from the CO_2_ anaesthesia for at least two days before measurements (MacMillan et al., 2017). All experiments were performed on 4-9 days-old non-virgin female flies, because of their larger size.

### Heat tolerance assays

Behavioural heat tolerance phenotypes were characterised with a ramping and a static assay using the same setup as previously described in Jørgensen *et al.* (2019). In this setup the fly was exposed to homogenous heat exposure within a glass vial that was submerged in a water tank with a controlled temperature (Fig. 1A). In the ramping assay, temperature was increased by 0.25 °C min−1 from 19 °C. Two behavioural phenotypes were recorded during this experiment: 1) the temperature at which the fly would lose coordination and fall on its back (T_back_) and 2) the temperature at which the fly was completely still (T_coma_). T_coma_ was verified by poking the vial lids with a stick to agitate the flies and check for reflexes. The static assay used a similar setup and method to record knockdown, but instead of increasing the temperature gradually, the flies were placed in the bath pre-set to 38 °C, after which the exposure durations causing loss of coordinated movement (t_back_) and heat coma (t_coma_) were noted (here the lowercase “t” represents time). The “static” assay was only static for 1 hour at 38 °C after which the temperature was increased by 0.25 °C min^−1^ to ensure that more heat tolerant flies would also succumb to heat stress. 7 flies were measured for each species in each assay, except *D. subobscura* in the ramping assay (n=6).

**Fig. 1.**
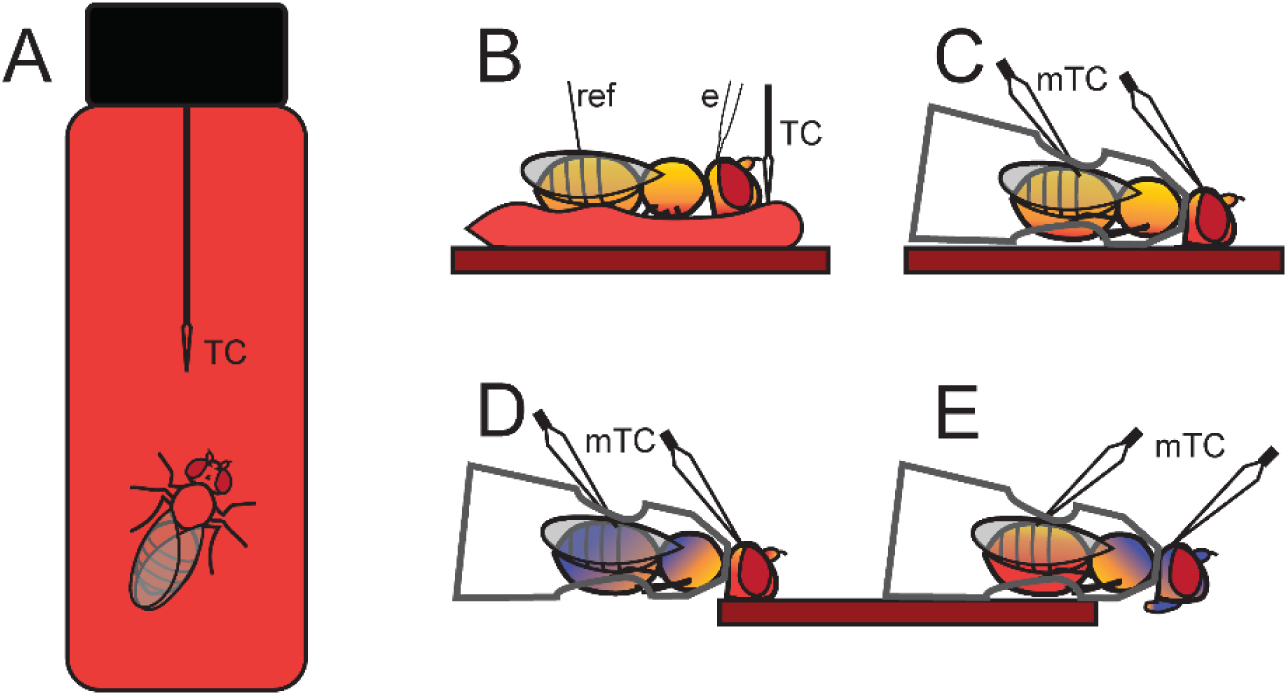
Overview of heating methods used for the experiments. Colour of the fly body indicates the assumed heat distribution, with red indicating warmer over yellow to blue for colder (eyes are red to characterise *Drosophila*). Spear-shaped arrows show the placement of thermocouples for each method, normal size (1.5 mm tip) thermocouples are marked with TC, micro thermocouples (25 μm tip) with mTC. (A) For behavioural phenotype assessment the fly was placed in a glass vial which was submerged in a temperature-controlled water bath. A uniform heat distribution around the fly was expected. (B) To measure spreading depolarisation the fly was fastened in a bed of wax (lighter red) on top of a Peltier element heating stage (darker red). The wax bed is assumed to give a relatively uniform heat distribution across the ventral body surface, but the dorsal side is possibly cooled slightly by the surrounding air. For these experiments, temperature was measured on top of the wax, adjacent to the head. The placement of the reference (ref) and measuring electrode (e) is also shown. (C) To assess heat sensitivity following whole-body heat exposure the fly was tethered inside a pipette tip, which was placed on the heating stage (dark red). The ventral side was warmer than the dorsal side, and the head tended to be slightly warmer than the abdomen. For these experiments we measured temperature on the dorsal side of the head and abdomen using micro-thermocouples. (D) In selective heating of the head, the fly was tethered but here only the head was in contact with the heating stage. Consequently, the abdomen and thorax were maintained at a lower temperature. (E) Selective heating of the abdomen resulted in a lower temperature of the thorax and head, notice that the non-measuring parts of thermocouples are oriented away from the heating plate.

### Measuring spreading depolarisation

Electrophysiological measurements of DC potentials in the CNS (a proxy for nervous function) were carried out as described by Andersen *et al*. (2018). Filamented borosilicate glass capillaries (1 mm diameter; 1B100-F-4, World Precision Instruments, Sarasota, Florida, USA) were pulled to low tip resistance (5-7 MΩ) using a Flaming-Brown PC-84 micro-pipette puller (Sutter Instruments, Novato, CA, USA) and back-filled with 500 mM KCl solution. The glass electrodes were connected to a Duo 773 intracellular differential amplifier (World Precision Instruments, Sarasota, Florida, USA) using the low impedance channel and probe, and a chlorinated Ag/AgCl wire was used as reference electrode to ground the preparation. An MP100 data-acquisition system was used to digitalize the voltage output which was recorded using AcqKnowledge software (Biopac Systems, Inc., CA, USA).

A fly was prepared for measurement by gently fastening its ventral side to a bed of wax on a glass cover slide. Using a small pair of scissors, a small hole was cut in the abdomen between the second and third-to-last tergites for placement of the ground electrode. Another cut was made along the head midline just posterior to the ocelli to insert the glass recording electrode. The cover slide with the fly was placed onto a Peltier plate pre-set to 30 °C which could be thermoelectrically heated (PE120, Linkam Scientific Instruments, Tadworth, United Kingdom), and temperature was monitored continuously using a type K thermocouple (integrated with the MP100 data-acquisition system) placed on top of the wax, adjacent to the head of the fly (Fig. 1B). This heating method was expected to heat the ventral side of the fly homogeneously, but also result in a small temperature gradient from the ventral to the dorsal side. The glass electrode and the reference (Ag/AgCl) electrode were placed in their designated holes using micromanipulators, and the voltage was zeroed. To test the quality of the preparation, a flow of humidified N_2_ was passed over the fly to elicit an anoxic spreading depolarisation (SD). The single depolarisation triggered by anoxia, persists throughout the exposure to N_2_, but has been found to be completely reversible in *Drosophila* (Armstrong et al., 2011; Rodríguez & Robertson, 2012) and locusts (Rodgers *et al*., 2007), and additionally we did not find any difference in timing of SD in heating experiments with and without prior anoxia treatment. We therefore used this anoxia test to discard preparations that failed to depolarise (suggesting that there was a problem with the electrode placement). This test also gave an indication of the size of depolarisation that could be expected from that particular preparation as this is also dependent on the quality of impalement and location of the recording electrode. If the preparation had depolarised ≥20 mV in response to anoxia, the voltage was zeroed again, and the preparation was either used for ramping, static or control experiments.

In ramping experiments, the temperature of the thermal stage was increased from 30 °C by 0.25 °C min^−1^ and the temperature (at the half-amplitude of the negative DC shift associated with SD) of the first and last SD event (SD_first_ and SD_last_, respectively) along with the number of SD events was recorded. The ramping continued until it was clear that no more depolarisations would occur, which was concluded when the preparation could no longer maintain a stable base line DC potential (see example traces in Fig. 2). In static heat exposure experiments, temperature was rapidly increased from 30 °C to 38 °C (mean heating time: 73 s, approx. 6.6 °C min^−1^), and the timing of SD_first_ and SD_last_ and the number of depolarisation events were noted as above. The stage was kept at 38 °C until no more depolarisations were anticipated (same criterion as in ramping experiments). In preparations for which no depolarisations had occurred during the 1-hour exposure (only in *D. melanogaster* and D. *mojavensis*), the stage temperature was increased by 0.25 °C min^−1^ after the first hour at 38 °C and this heating was continued until depolarisations were measured. Some of the preparations elicited only a single SD event, and accordingly the temperature/time reported was the same for SD_last_ as SD_first_ (see Fig. 2C).

**Fig. 2.**
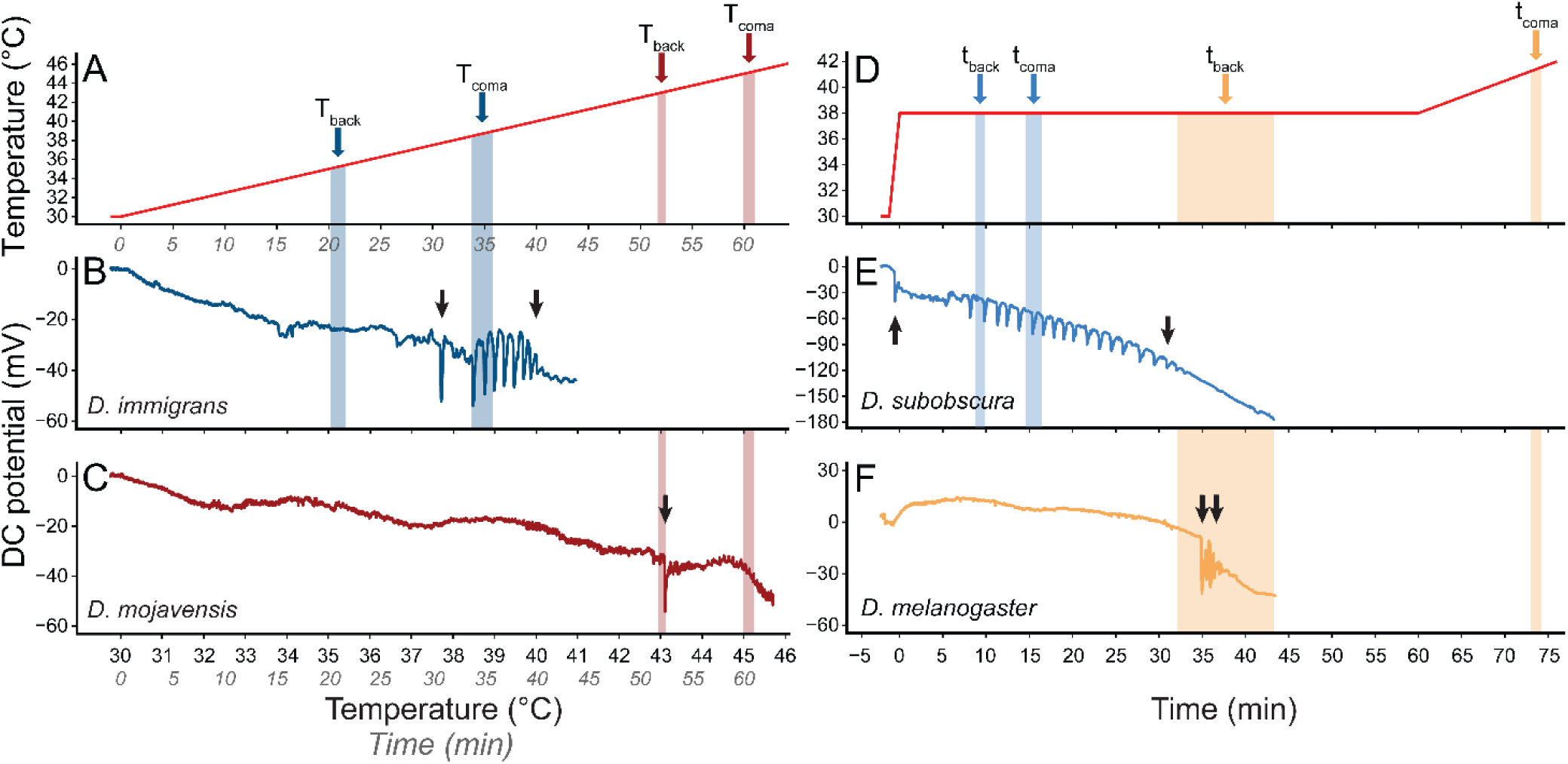
Representative temperature and DC potential traces from ramping (A-C) and static (D-F) heat exposures. The temperature profiles during (A) ramping and (D) static assays are marked with species-coloured arrows and transparent boxes for the two phenotypes, T/t_back_ and T/t_coma_ (mean ± s.e.m.), for two species from each assay (The phenotypes and DC potential traces were not recorded from the same individuals). (B) The heat sensitive *D. immigrans* experienced spreading depolarisation at a lower temperature than the (C) heat tolerant *D. mojavensis* during a ramping assay. (E) Similarly, the heat sensitive *D. subobscura* experienced spreading depolarisation sooner than the (F) more heat tolerant *D. melanogaster* in the static assays. In (A-C), the x-axis show both measured temperature and the corresponding time (italicised) according to the ramping rate of 0.25 °C min^−1^, and in (D-F) the time scale is adjusted such that time = 0 when the temperature reached 38 °C. Black arrows in (B-C, E-F) mark the SD_first_ (left) and SD_last_ (right) SD event, notice the example of a single SD event in *D. mojavensis* (C).

A number of pilot studies were conducted to test if the starting condition at 30 °C or the handling of the fly was stressful enough to elicit SDs by keeping a few *D. immigrans* (the least heat tolerant species) and *D. mojavensis* (the most heat tolerant species) at 30 °C for 1 hour, but these conditions failed to elicit SDs in either species. These experiments were concluded by increasing temperature by 1 °C min^−1^ until SD events were observed, leading us to conclude that the preparations were responsive but that the handling and starting conditions (30 °C) alone were unable to evoke this response.

### Selective heating of head and abdomen

To further examine the role of nervous function in heat tolerance, we performed a series of experiments in which we selectively heated the head or the abdomen of flies and compared their survival after 24 hours to that of flies that had been heated more uniformly (See Fig. 1C-E). The motivation for this study was to examine if the head (dominated by nervous tissue) was more heat sensitive than the abdomen (dominated by fat-body and intestinal tissue). Only three species (*D. subobscura, D. melanogaster* and *D. mojavensis*) were used for these experiments as they represent low, medium and high heat tolerance, respectively. *D. subobscura* was chosen to represent low heat tolerance rather than *D. immigrans* due to its smaller size, which made it more appropriate for the method.

For these experiments the flies needed to be restrained in a way that allowed one end of the fly to be held closer to the heating stage, and as survival was used as the measure of sensitivity, the restraining method fixation should also allow for the flies to be moved from the heating stage without inflicting injury to the animals. Accordingly, flies were fastened in 200 μL pipette tips, using a device originally designed for hemolymph extraction (MacMillan & Hughson, 2014). With a stream of air, the fly was manipulated headfirst into the pipette tip, and the airflow was blocked once the fly was stuck in the tip (taking care not to injure it). The pipette tip was removed from the device and the tip was cut off just anterior to the head followed by two cuts (one from the dorsal and one from the ventral view of the fly) that were made in roughly a 45°C degree angle towards the anterior part of tip (Fig. 1C-E). These angled cuts allowed better contact between the head and the heating stage on the ventral side and room for the thermocouple to measure head temperature on the dorsal side. Using a scalpel, some of the plastic covering the abdomen was gently “shaved” off, while making sure that no holes were made. The tip was then reattached to the air pressure device and the fly was “pushed” until the head protruded from the tip. The area that had been thinned before was now cut away, leaving the abdomen exposed, thereby decreasing the distance to the heating stage on the ventral side (Fig. 1C-E). Another cut was made in the dorsal side of the tip allowing placement of a micro thermocouple directly on the dorsal side of the abdomen (here it was often necessary to move the wings to the side) (Fig. 1C-E). Flies that were injured (other than severed wings) were discarded. The preparations were used for either whole-body heating, selective heating of the head, selective heating of the abdomen or as un-heated controls. Flies were generally heated on the ventral side, but we also tested some flies exposed to whole body heating from the dorsal side (see Supplements Fig. S1).

For ventral whole-body heating, the pipette tip was placed on the Peltier plate (PE120, Linkam Scientific Instruments, Tadworth, United Kingdom) with the wide end of the tip at a slightly positive angle, to facilitate closer contact between the heating stage and the ventral side of the head and abdomen (Fig. 1C). When the tip was staged, two micro K type Fine thermocouples (tip diameter 25μm, KFG-25-100-100, ANBE, Genk, Belgium) were placed on the surface of the head and the abdomen, respectively (Fig. 1C). This method gave a relatively homogenous heating of the fly when measured on the dorsal side, with a tendency for slightly higher temperatures measured on the head (possibly due to closer contact with Peltier plate). For every sample, the tip was turned 180° horizontally, such that the head and abdomen switched location on the heating stage, to minimise any differences in heating across the stage. The transversal temperature gradient that arose from ventral heating was measured in *D. mojavensis* by gradually moving thermocouples through head and abdomen from the dorsal towards the ventral side, in flies that had been killed before the experiment. This transverse difference was recorded at 2.51 ± 0.22 °C and did not differ between head and abdomen (one sample t-test, t=11.05, *df*=11, *p*<0.001). Similar measurements were made for a few *D. melanogaster* and *D. subobscura*, with comparable results.

To test heat tolerance, the temperature of the heating stage was quickly increased to the desired test temperature (∼1.5 min), and once the temperature was stable the fly was left at this condition for 15 minutes. After heating, temperature would rapidly drop to room temperature (∼1 min) when the thermal stage was turned off. The tip was then removed from the Peltier plate, and the fly was immediately checked for movement. After 15 minutes, the fly was again checked for movement, released by cutting the tip and then transferred to a 2-mL Eppendorf tube with fly medium in the bottom and air holes in the lid. Flies were checked for movement after one day of recovery following the heat exposure (recovery at 19 °C), and their status (live/dead) here was used for further analysis. Flies were regarded as “dead” if they were unable to move after the 24-hour recovery period.

Selective heating of either head (Fig. 1D) or abdomen (Fig. 1E) was performed using the same preparation as above, but with the body part to be heated placed on the heating stage while the rest of the body was placed away from the stage. This heating method resulted in large temperature differences between body parts, with heating of the head giving a larger difference than heating of the abdomen (Table 1).

**Table 1.**
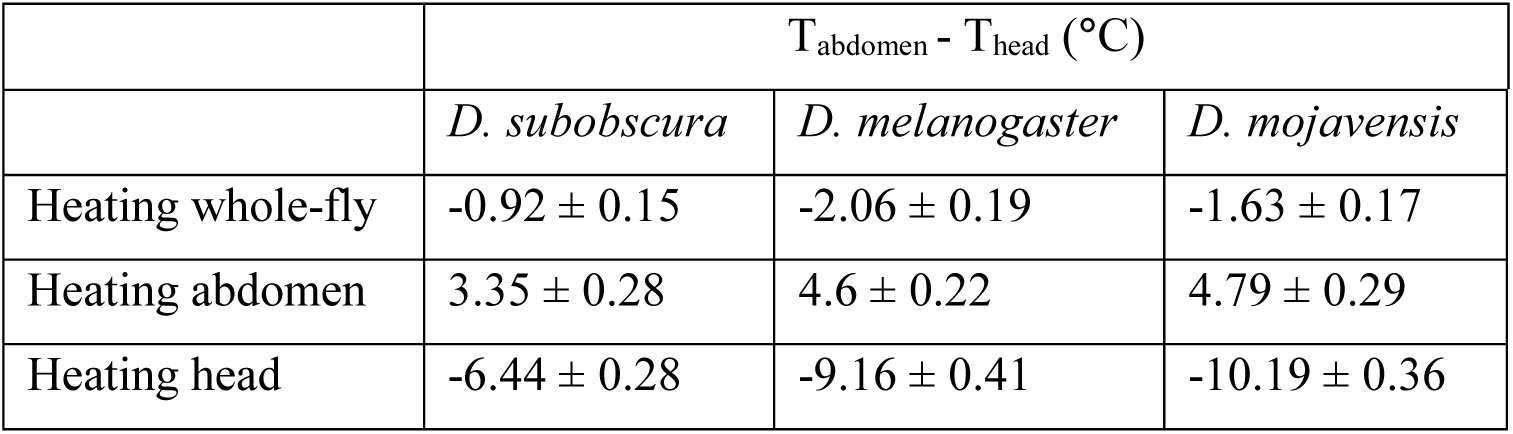
Temperature difference between abdomen and head measured topically on the dorsal side with ventral heating. Values reported as mean ± s.e.m.

|  | T<sub>abdomen</sub> - T<sub>head</sub> (°C) |  |  |
| --- | --- | --- | --- |
|  | *D. subobscura* | *D. melanogaster* | *D. mojavensis* |
| Heating whole-fly | -0.92 ± 0.15 | -2.06 ± 0.19 | -1.63 ± 0.17 |
| Heating abdomen | 3.35 ± 0.28 | 4.6 ± 0.22 | 4.79 ± 0.29 |
| Heating head | -6.44 ± 0.28 | -9.16 ± 0.41 | -10.19 ± 0.36 |

Control experiments were performed to test if the manipulation of the flies resulted in any mortality. In these experiments, the flies were prepared similarly to flies used for heating, but instead of heat exposure they were kept at room temperature and assessed for survival following the same protocol.

### Data analysis

All data analyses were performed in R version 3.5.2 (R Core Team, 2018). Unless otherwise stated all results are reported as mean ± s.e.m., and the critical value for statistical significance was 0.05. Onset of the phenotypes (T_back_ and T_coma_) and SD events (SD_first_ and SD_last_) were tested for co-occurrence using two-way ANOVAs for each assay type (ramp and static) with species and measured variable (T_back_, T_coma_, SD_first_, SD_last_) in ramp and (t_back_, t_coma_, SD_first_, SD_last_) in static assays as factor variables. Tukey’s HSD *post hoc* test was used to examine differences in onset of phenotypes and SD events within species. The correlation between heat stress phenotypes and onset of SD events was examined *between* species *within* assay type using linear regressions (lm()-function in R). The regression lines were compared to the line of unity (intercept = 0, slope =1) with the function linearHypothesis in the *Car*-package (Fox & Weisberg, 2011).

The survival assessments from the selective and whole-body heating experiments were paired with the temperatures measured from the thermocouples placed on head and abdomen. The temperature causing 50% mortality (LT_50_) after 24 hours was estimated through a non-linear least square-model using the nls()-function in R. The nls()-function was given the following equation of a sigmoidal curve: 

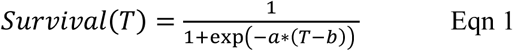

Where *Survival(T)* is survival at the temperature *T, a* is the slope of the descending part of the sigmoidal curve and *b* is the estimate of LT_50_. 95% level confidence intervals were calculated for each survival curve around the estimated LT_50_ using confint2() from the *nlstools*-package (Baty *et al*., 2015). Curves with non-overlapping confidence intervals were regarded significantly different.

## Results

### Loss of CNS function and onset of heat stress phenotypes

Neural function during heat exposure was examined by measuring negative DC shifts associated with spreading depolarisation (SD) in the central nervous system (CNS) in the head of five *Drosophila* species representing a range of heat tolerances. Flies were heated using either a ramping assay during which temperature (i.e. stress intensity) was gradually increased, or a static assay during which temperature was kept constant at 38 °C. The temperature (ramp) or time (static) of the first or last SD (SD_first_ and SD_last_, respectively) were then compared to the timing or temperature of two behavioural heat stress phenotypes measured using similar heating protocols (the phenotypes measured were the loss of coordinated movement (T/t_back_) and onset of heat coma (T/t_coma_), Fig. 2). These experiments were used to examine 1) if heat stress phenotypes correlate with signs of neural dysfunction, and 2) if this putative correlation is affected by the way heat stress is inflicted.

When flies were exposed to gradually increasing temperatures in a ramp, there were clear interspecific differences in the temperatures where the behavioural heat stress phenotypes were observed. For example, the least heat tolerant species (*D. immigrans*) showed loss of coordination (T_back_) at 35.22 ± 0.45 °C and went into heat coma (T_coma_) at 38.69 ± 0.25 °C, while the most heat tolerant species (*D. mojavensis*) reached T_back_ at 43.01 ± 0.24 °C and T_coma_ at 45.11 ± 0.34 °C, giving the species system a range of T_back_ of 7.8 °C and T_coma_ of 6.4 °C. Similarly, the temperatures at which SD events were observed gave interspecific differences of 7.4 °C for SD_first_ and 6.5 °C for SD_last_ between the least and most tolerant species (again *D. immigrans* and *D. mojavensis*). Generally, we found that the temperature of T_back_ and T_coma_ coincided with perturbation of nervous function as indicated by SD_first_ and SD_last_ (Fig. 3). For three of the species (*D. mercatorum, D. melanogaster* and *D. mojavensis*) the two-way ANOVA followed by a Tukey HSD *post hoc* test did not reveal any significant differences in temperature between either of the behavioural phenotypes and the SD events. For the remaining two species (also the two least tolerant), T_coma_ was observed at a significantly higher temperature than the first SD event (Fig. 3). In *D. immigrans* it was also possible to separate the two heat stress phenotypes from each other, as T_back_ was observed at a significantly lower temperature than T_coma_. However, we caution that the means of heating differed between the phenotype experiments and the neurological experiments, and that this could be a source of experimental noise (see Methods and Discussion for further arguments). To test if there was a general co-occurrence of phenotypic and neurological events, we performed linear regressions of the mean temperatures of either of the two behavioural phenotypes and the two neuronal phenotypes (Table 2). All regression combinations yielded high coefficients of determination (*R*^*2*^: 0.73-0.9), and only one of the four regressions (SD_first_ against T_coma_) was significantly different from the line of unity (Table 2, see Supplements Fig. S2). The regression analysis indicated that across species there were generally only small differences between the temperature where behavioural and neurological collapse was observed.

**Table 2.** Coefficients of determination (*R*^*2*^) from linear regressions between behavioural phenotypes and SD measurements, *p*-values are from the test comparing the linear regressions to the line of unity (i.e. *p*-values above 0.05 indicate that the compared phenotypes occur at the same temperature/time). The highest *R*^*2*^ in each assay type is marked in bold italics, and linear regressions which were different from the line of unity (*p* < 0.05) are underlined. See Supplements Fig. S2-S3 for a graphical representation of the linear regressions.

|  | Dynamic |  | Static |  |
| --- | --- | --- | --- | --- |
|  | T<sub>back</sub> (°C) | T<sub>coma</sub> (°C) | t<sub>back</sub> (min) | t<sub>coma</sub> (min) |
| SD<sub>first</sub> (°C or min) | $R^2 = 0.73, p = 0.699$ | $R^2 = 0.8, p = \underline{0.041}$ | $R^2 = \mathbf{0.86}, p = 0.812$ | $R^2 = 0.65, p = 0.309$ |
| SD<sub>last</sub> (°C or min) | $R^2 = 0.78, p = 0.260$ | $R^2 = \mathbf{0.9}, p = 0.089$ | $R^2 = 0.77, p = 0.392$ | $R^2 = 0.65, p = 0.632$ |

**Fig. 3.**
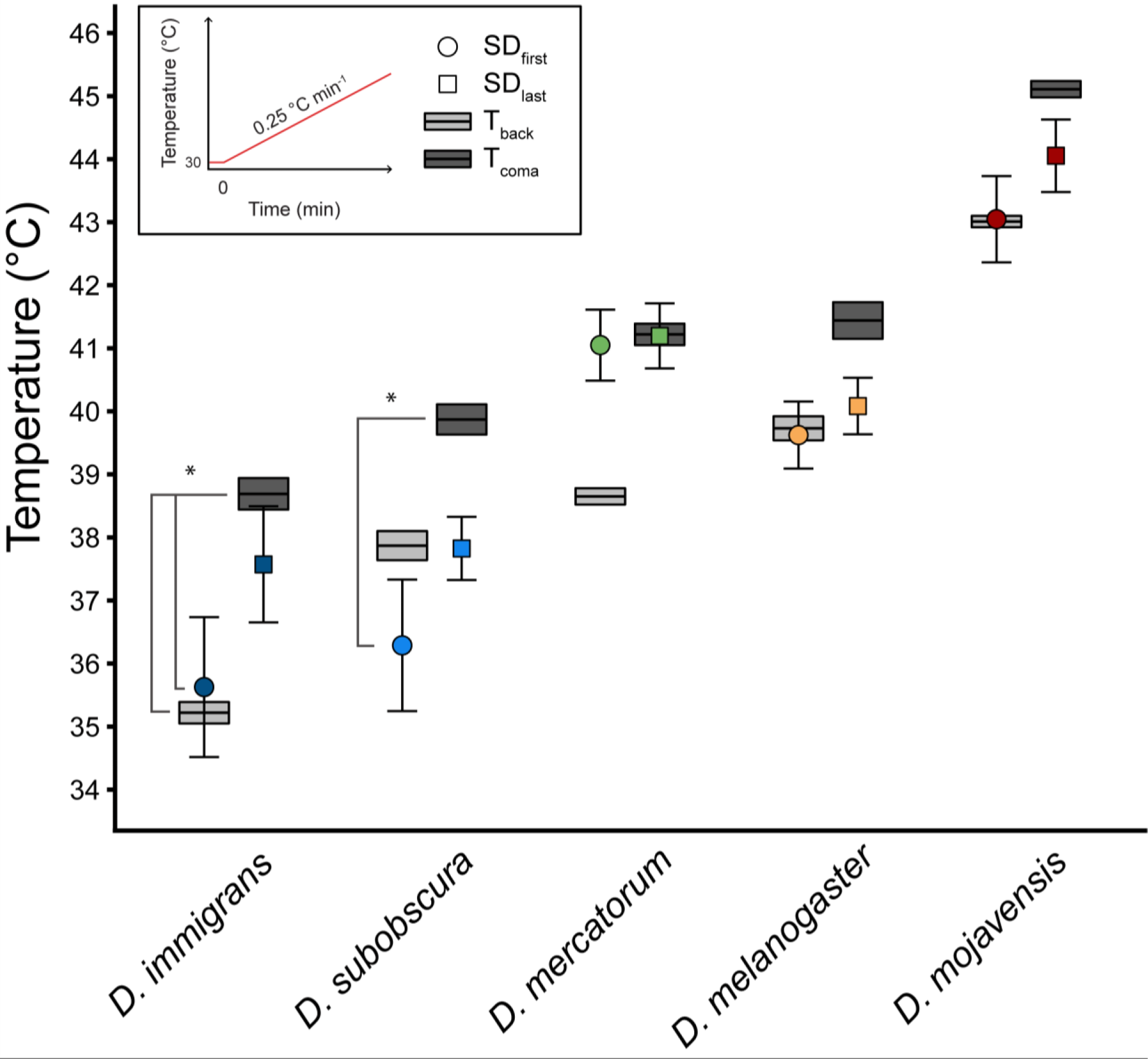
Temperature of SD_first_ (circle) and SD_last_ (square) and the temperature of the two behavioural heat stress phenotypes T_back_ (light grey bars) and T_coma_ (dark grey bars) in a ramping assay. SD_last_ coincide with SD_first_ in cases where only a single SD event was observed. SD measurements were performed on a Peltier element while the whole animal knockdown phenotype were observed from flies in glass vials submerged in a temperature-controlled water bath. Asterisks mark significant differences between either of the four phenotypes (*p*<0.05), n=7 for each species and data are reported as mean ± s.e.m.

During constant heat exposure (38 °C, Fig. 4), we recorded the timing of SD events and behavioural heat stress phenotypes and again we found these behavioural and neurological measures to coincide. Note that for some species we started to increase the temperature by 0.25 °C min^−1^ after 1 hour of exposure, but that all measures are reported in minutes of exposure. Between species there was a clear increase in the heat exposure duration that the nervous system could uphold function with increasing heat tolerance of the species (according to the timing of behavioural heat stress phenotype onset), although the least tolerant species in terms of neuronal failure (*D. subobscura*) was the second least tolerant when assessed for behavioural phenotype (*D. immigrans* was the least tolerant on this term, as in the ramping assay) (Fig. 4). A two-way ANOVA followed by a Tukey HSD *post hoc* test revealed that it was not possible to separate the timing of behavioural heat stress phenotypes and the neurological perturbations in *D. immigrans, D. subobscura* and *D. mojavensis*. In *D. mercatorum* and *D. melanogaster* significant differences between the timing of behavioural and neurological phenotypes were found, with a delayed coma onset for *D. melanogaster* relative to both t_back_ and the SD events, and a relatively long time span between the loss of coordinated movement and the last SD event in *D. mercatorum* (Fig. 4). However, linear regressions on the mean time of the four possible combinations of SD events and behavioural phenotypes showed a high correlation between both SD_first_ and SD_last_ with t_back_ (*R*^*2*^: 0.77-0.86), while the correlations between SD types and t_coma_ were slightly weaker (*R*^*2*^: 0.65) (Table 2, see Supplements Fig. S3). When the four regression lines were compared to the line of unity, none of them were significantly different, again suggesting that across the species system there were generally an overlap between the exposure durations that resulted in behavioural and neurological phenotypes.

**Fig. 4.**
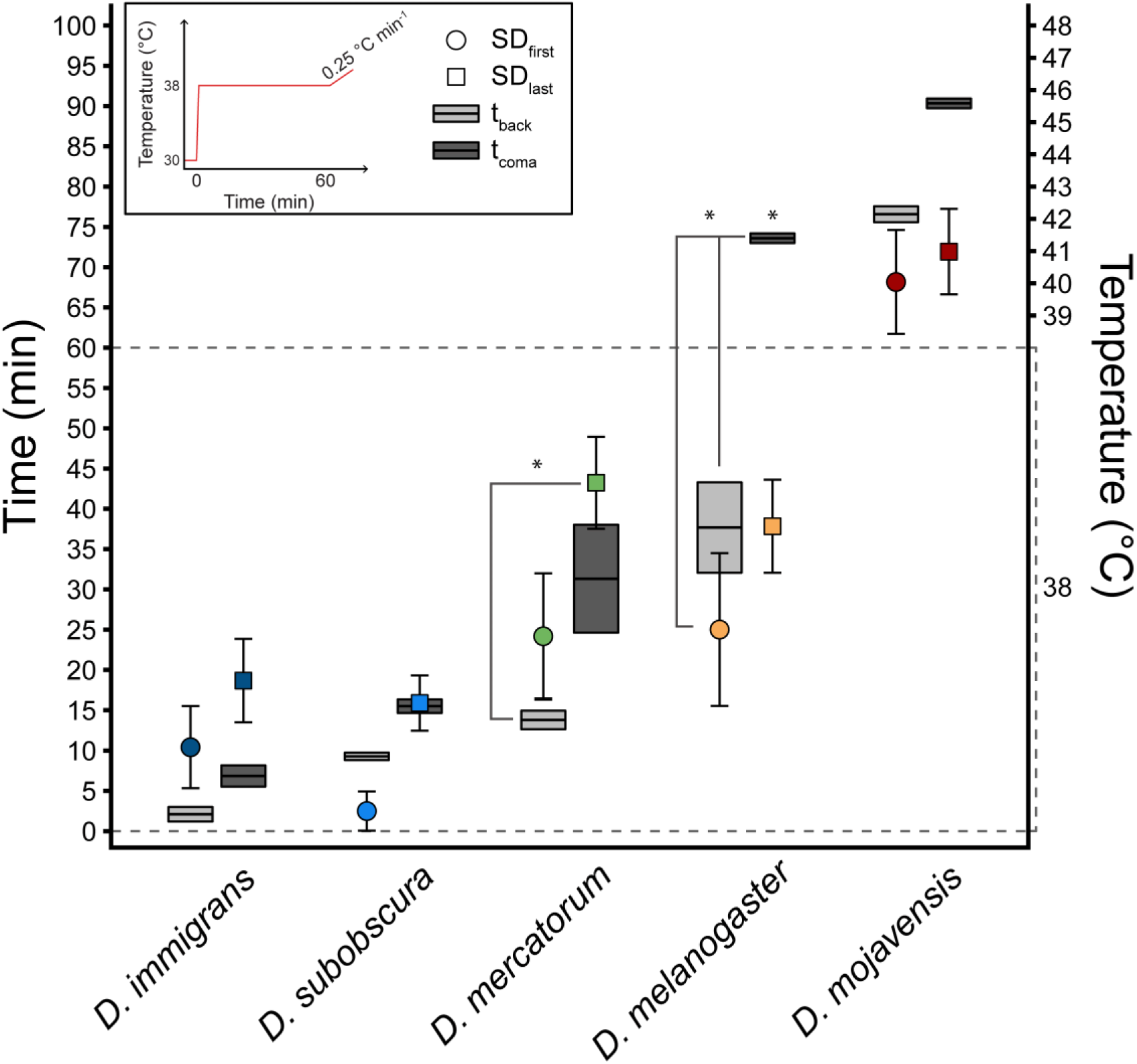
Exposure time in a static assay until SD_first_ (circle) and SD_last_ (square) and loss of coordinated movement (t_back_, light grey bars) and onset of coma (t_coma_, dark grey bars). The time scale is adjusted such that time = 0 when the temperature reached 38 °C (average time to heat from room temperature to 38 °C was 73 s for SD measurements). After 1 hour at 38°C the temperature was increased by 0.25 °C min^−1^, and SDs and phenotypes that occurred during the ramp is here presented on the time scale (with the corresponding temperature on the secondary y-axis). SD measurements were performed on a Peltier plate while behavioural phenotypes were assessed from flies in glass vials submerged in a temperature-controlled water bath. Asterisks mark significant differences between either of the four phenotypes (*p*<0.05), n=7 for each species and data are presented as mean ± s.e.m.

Examination of the DC potential measurements showed considerable variance between preparations. Some preparations where characterised by only eliciting a single SD event (meaning that SD_first_ and SD_last_ occurred at the same time/temperature, Fig. 2C) while other specimens showed multiple (2-30) SD events (see examples in Fig. 2). Comparison between the ramping and constant heat exposures showed that single SD events were much more prevalent during the ramping heat exposure (40% of individuals showed single SD, n=35) than in the constant heat exposure (9% showed single SD, n=29) (see Supplements Fig. S4). Furthermore, when the constant heat exposure for 1 hour was followed by a ramping increase in temperature, flies would mostly elicit just a single SD (66%, n=6). All five species were able to display both single and repeated SD events and in roughly the same proportion (2-4 preparations of each species (out of 7) showed a single SD during ramping). The number of SD events observed in “multiple” SD events also differed with heat exposure assay. In static assays, preparations with multiple SDs elicited 11.38 ± 1.56 SD events while preparations with multiple SDs during ramping assays only had 5.95 ± 1.12 SD events (two sample t-test, t=2.83, *df*=43.15, *p*=0.007).

### Selective heating of the head and abdomen

As heat coma and heat death often occur in close succession, we performed an experiment designed to investigate and compare the heat sensitivity of the head (site of nervous function measurements from the first experiment) and the abdomen (consisting more of visceral tissues) (see Fig. 1C-E). This test involved restraining flies in pipette tips and non-heated controls for handling showed 0% mortality for *D. subobscura* and *D. melanogaster*, and 13% mortality for *D. mojavensis* after 24 hours (n=14/16/39, respectively). For these experiments the temperature estimated to cause 50% mortality in the flies 24 hours after heat exposure (LT_50_) was used to compare heat sensitivity between body parts.

Both whole-fly and selective heating showed that the heat tolerant *D. mojavensis* had higher values of LT_50_ than the moderate heat tolerant *D. melanogaster*, which in turn also had higher values of LT_50_ than the heat sensitive *D. subobscura* (Fig. 5). When the whole fly was heated simultaneously, we did record differences between head and abdominal temperature (measured topically on the dorsal side), but these differences were generally less than 2 °C (see Table 1 and Supplements Fig. S1). In experiments using selective heating of either the head or abdomen the flies were characterised by much larger regional differences in temperature (ΔT ranging 3.35-10.19 °C depending on species and body part heated, see Table 1).

**Fig. 5.**
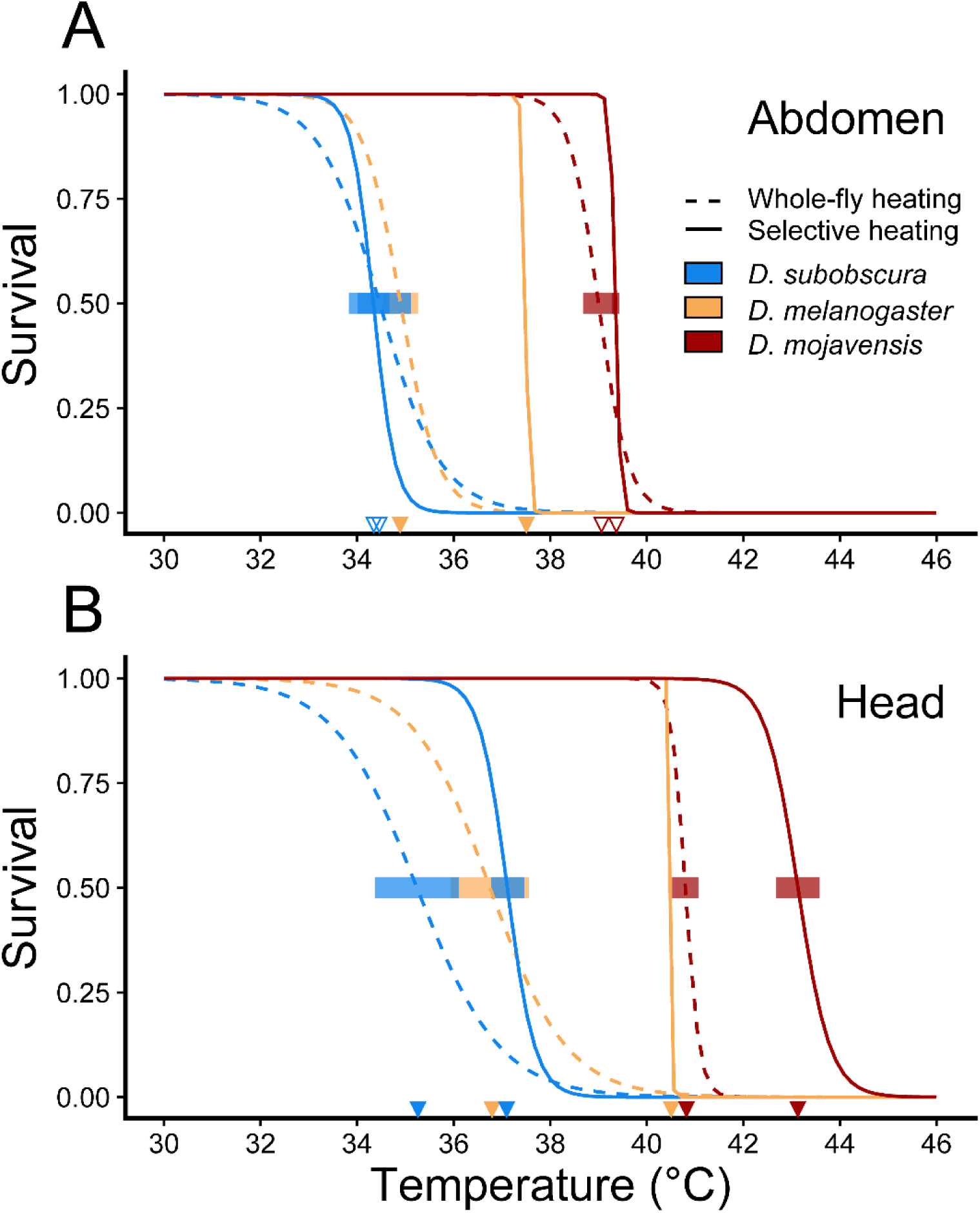
Survival curves and LT_50_ estimates for whole-fly and selective heating of *D. subobscura* (blue), *D. melanogaster* (yellow) and *D. mojavensis* (red). (A) Survival curves are related to the temperature measured topically on the abdomen during selective heating of the abdomen (full lines) and whole-fly heating (dashed lines). (B) Survival curves are related to the temperature measured topically on the head during selective heating of the head (full lines) and whole-fly heating (dashed lines). *Note that whole-fly heating curves are slightly different in A and B because they are based on the temperature measurements from the abdomen and head, respectively.* LT_50_, the temperature that resulted in 50% mortality, was estimated for all survival curves, and is marked on the temperature axis by a species coloured triangle. If the 95% confidence intervals of selective heating and whole-fly heating LT_50_ (shaded, species coloured areas) within a species did not overlap, a closed triangle was used, and conversely, if confidence intervals overlapped, open triangles were used. Whole-fly heating and selective heating of abdomen and head were performed on n=24/15/18 for *D. subobscura*, n=24/17/16 for *D. melanogaster* and n=35/17/17 for *D. mojavensis*, respectively. Selective heating of *D. melanogaster* yielded very steep survival curves where the confidence intervals could not be determined.

The experiments revealed species specific differences in the relation between LT_50_ estimates during whole animal heating and selective heating. For *D. mojavensis*, heating the abdomen (and maintaining the head at a lower temperature, ΔT=4.79 ± 0.29 °C) did not change the LT_50_ compared to abdominal temperature when the whole fly was heated (LT_50_ was 0.35 °C higher but the estimates have overlapping 95% confidence intervals, Fig. 5A). Thus for *D. mojavensis*, LT_50_ was the same irrespective if the head was kept cool or warm during heating of the abdomen. When the head of *D. mojavensis* was heated selectively (with the abdomen considerably cooler: ΔT=10.19 ± 0.36 °C), LT_50_ increased by 2.33 °C compared to flies experiencing whole animal heating (non-overlapping 95% confidence interval, Fig. 5B). Thus, a higher head temperature was needed to evoke mortality in *D. mojavensis* when the abdomen was relieved from heat stress.

Performing the experiments on *D. melanogaster* we observed slightly smaller differences between body parts than in *D. mojavensis*, both when the head was selectively heated (abdomen maintained at a lower temperature, ΔT=9.16 ± 0.41 °C) and when the abdomen was heated (head kept cooler, ΔT=4.6 ± 0.22 °C). For *D. melanogaster* we found LT_50_ to increase when applying selective heating on the abdomen (LT_50_ was 2.59 °C higher, Fig. 5A) and the head (LT_50_ was 3.77 °C higher, Fig. 5B), compared to LT_50_ resulting from whole-fly heating. Accordingly, maintaining one end of a *D. melanogaster* at a lower temperature than the other, increases heat tolerance of the fly.

In experiments with *D. subobscura*, the temperature differences between body parts were smaller than for the other two species. Selectively heating the abdomen made the abdomen 3.35 ± 0.28 °C warmer than the head but did not change the LT_50_ of the abdomen when compared to that of whole-fly heating (LT_50_ was 0.13 °C lower for the selective heating, likely attributed to the shape of the survival curve, but with overlapping 95% confidence intervals). When selectively heating the head, resulting in a 6.44 ± 0.28 °C colder abdomen, head LT_50_ increased by 1.87 °C compared to head LT_50_ of whole-animal heated flies.

## Discussion

Inter- and intraspecific differences in heat tolerance have been demonstrated for *Drosophila* in multiple studies (Castañeda et al., 2015; Jørgensen et al., 2019; Kellermann et al., 2012; Kimura, 2004; Overgaard et al., 2014; Stratman & Markow, 1998). These differences have often been measured using the onset of reversible behavioural phenotypes such as loss of coordinated movement and entry into heat coma, or by measuring heat induced mortality in animals exposed to high temperatures (Lutterschmidt & Hutchison, 1997a). However, it is still unclear which physiological perturbations are the proximate cause of the different heat tolerance endpoints (but see Robertson (2004) and Rodgers *et al*. (2010)), and this has been particularly difficult to discern because of the close proximity of the endpoints at high temperatures. Multiple physiological mechanisms have been suggested as the proximate cause of heat mortality, including oxygen transport limitations, protein denaturation, loss of membrane integrity or ion homeostasis, and mitochondrial dysfunction (Bowler, 2018; Davison & Bowler, 1971; Gladwell, 1975; Pörtner, 2001; Somero, 1995). The endpoint prior to mortality, the onset of heat coma, has instead been suggested to be caused by either muscular or nervous failure (Bowler, 1963; Gladwell *et al*., 1975; Robertson, 2004). In locusts exposed to increasing temperature, ventilation failed concurrently with an abrupt surge in extracellular [K^+^], which has been related to a drop in DC potential that is a reliable marker of spreading depolarisation in the CNS (SD) (Robertson, 2004; Rodgers *et al*., 2007). Once the locust was returned to benign temperatures, extracellular [K^+^] surrounding the neurons returned to baseline levels, and the motor pattern ventilation resumed (Rodgers *et al*., 2007; Rodgers *et al*., 2010).

To our knowledge there has been no comprehensive comparative studies investigating species differences in CNS function at high temperature and the aim of this study was to examine the role of the nervous system in relation to heat tolerance in five *Drosophila* species. The temperatures at which two behavioural phenotypes (loss of motor control (T_back_) and loss of motor function (T_coma_)) were observed were compared to the temperature of neuronal failure (SD) as assessed by electrophysiological measurements of DC potentials in the fly brain during ramping heat exposure, and likewise the timing of SD and behavioural phenotypes during constant heat exposure. These experiments revealed a good correlation between the failure of motor control/function and neuronal failure, however it is unclear if failure of the CNS is also causing heat mortality. Thus, we designed an experiment to test the sensitivity to heat exposure on different parts of the fly body to further examine if the nervous system could be limiting heat stress survival.

### Heat stress phenotypes correlate with onset of nervous failure

Measurements of spreading depolarisation (i.e. large negative shifts in DC potential) during both ramping and static assays, showed that, overall, perturbation of nervous function correlated well with the two behavioural heat stress phenotypes (t/T_back_ and t/T_coma_) (Fig. 3-4). Onset times and temperatures of the behavioural coma phenotype were similar to the values previously reported in the five species measured in similar heat tolerance assays (Jørgensen *et al*., 2019). The loss of motor function was assessed on untethered flies in glass vials with a homogeneous temperature, whereas SD measurements required the flies to be fastened and furthermore a hole was cut in the head and abdomen to insert measurement electrodes (Fig. 1). The invasive preparation required for SD measurements could potentially alter heat tolerance, and we also observed a surprisingly large internal thermal gradient in the fly (sometimes more than 2 °C) when using the Peltier plate for heating. The differences in experimental protocols between behavioral and neurological experiments are likely to introduce some noise in the comparison between these experiments, particularly because we know already that the rate of heat injury accelerates extremely quickly at high temperature (Q_10_ of heat injury accumulation rate is often >10.000). Thus, very small differences in exposure temperature (or time) can separate tolerance and death during heat exposure (Jørgensen *et al*., 2019). Considering these sources of variation, it would be unexpected to find a perfect correlation between the two experiment types. Despite these “experimental challenges” we found clear patterns of association between loss of motor control and the occurrence of SD events in the CNS (Figs. 3 and 4).

Generally, the characteristics of heat stress phenotypes follow a progressive loss of motor control, from first hyperactivity, through loss of coordinated movement and spasms to the onset of heat coma or heat stupor where the animal is unresponsive (Cossins & Bowler, 1987; Heath & Wilkin, 1970; Lutterschmidt & Hutchison, 1997a). Accordingly, for these experiments it follows that the two behavioral phenotypes t/T_back_ and t/T_coma_ are bound in a way such that t/T_back_ will occur prior to (or at a lower temperature) compared to t/T_coma_. Similarly, the first SD must precede the last SD, unless only a single SD event is observed (in which case the first and last SD are the same). It is therefore tempting to conclude that SD_first_ is linked to t/T_back_ and likewise SD_last_ to t/T_coma_ but with the lack of clear statistical support for this, we will only conclude that it is likely that the two closely occurring behavioural phenotypes (t/T_back_ and t/T_coma_) are linked to the simultaneously occurring SD events (SD_first_ and SD_last_, respectively). The relation between behavioural phenotypes and nervous dysfunction has also been examined at low temperatures in different species of *Drosophila*, where temperature of cold coma onset is also highly correlated with the temperature of SD in the CNS of *Drosophila* (Andersen & Overgaard, 2019; Andersen *et al*., 2018). However, similar to our heat experiments it is difficult to determine specifically how first and last SD events are linked to loss of motor control (T_back_) or loss of movement (T_coma_). Importantly, there is no association between cold-induced SD events and cold mortality as insects can survive cold in a “comatose” state for long periods of time (MacMillan & Sinclair, 2011; Overgaard & MacMillan, 2017).

The present study found that single SD events (instead of multiple events) were more prevalent in ramping experiments than during static heat exposure (Supplements Fig. S4). Additionally, the number of SD events that occurred in preparations with more than one SD, was significantly higher during ramping heat exposure compared to static. In hyperthermic locusts single continuous SD events that persist until the heat exposure is removed are the most prevalent, but repetitive SD events have been observed in locusts treated with ouabain (Rodgers *et al*., 2009; Spong *et al*., 2014) and in hyperthermic brain slices from immature rats (Wu & Fisher, 2000). Contrary to hyperthermia, which is thought to lead to accumulation of [K^+^], ouabain is limiting K^+^ clearance through its inhibition of the Na^+^/K^+^-ATPase (Rodgers *et al*., 2009). According to Rodgers *et al*. (2009) the repetitive SD events are caused by transient surges in extracellular [K^+^] that are resulting from imbalances between accumulation and clearance of K^+^. A speculative explanation for the increased prevalence of single SD events in ramps could be that when temperature is gradually increased, the mitigation of the physiological conditions resulting in SDs (high extracellular [K^+^] in the space surrounding the CNS) cannot keep up as heat stress increases exponentially (Jørgensen *et al*., 2019), resulting in a total silencing of the CNS. Conversely, the static exposure may allow the fly to remove some of the [K^+^] that has accumulated in the extracellular space. This could relieve the condition causing the SD event and temporarily restore some nervous function until a new SD events occurs when K^+^ clearance is surpassed by the accumulation (Rodgers *et al*., 2010). Despite differences in experimental protocols we here clearly demonstrate that SD events in the CNS and the loss of motor function or entry into coma coincide in *Drosophila* species with different levels of heat tolerance. This indicates that loss of CNS function is the proximal cause to the onset of heat coma (CT_max_), a behavioural phenotype that is commonly used to describe animal heat tolerance. However, as found in cold *Drosophila*, it is also important to emphasise that the significance of nervous dysfunction in the onset of coma does not necessarily mean that the loss of nervous function directly results in heat death.

### Selective heating of the head and abdomen suggests interspecific differences in body part heat sensitivity

To investigate the role of the CNS failure for heat mortality, we designed an experiment to estimate heat sensitivity of the head and the abdomen when either the whole fly was heated, or when one body part was selectively exposed to a higher temperature than the rest of the fly. If CNS failure at high temperatures is the main cause of heat mortality, then we would expect that maintaining the head at a lower temperature than the abdomen should also lower mortality. Conversely, if the head was heated selectively, we would expect mortality to occur at the same temperature as when the whole fly was heated. Manipulations of body compartment temperatures have previously been used successfully in crayfish (Bowler, 1963), goldfish (Friedlander *et al*., 1976) and Atlantic cod (Jutfelt *et al*., 2019) to investigate the heat sensitivity of either heat coma or heat mortality. To our knowledge this is the first study to attempt such a study in small insects such as *Drosophila*.

Using the experimental setup with a fly tethered in a pipette tip, we found clear differences in heat tolerance (measured as LT_50_) between species, such that the desert species *D. mojavensis* was more heat tolerant than the cosmopolitan *D. melanogaster*, which in turn was more heat tolerant than the temperate *D. subobscura*. This finding is entirely consistent with the other heat stress phenotypes measured in the present study and with findings from previous studies (Jørgensen *et al*., 2019; Kellermann *et al*., 2012). The tethering of the flies was not in itself invasive as attested by no mortality of controls in *D. subobscura* and *D. melanogaster*, and low mortality in *D. mojavensis* controls. Selective heating of abdomen and head suggests interspecific differences in body part sensitivity (Fig. 5). All three species showed increased heat tolerance of the head when the abdomen was simultaneously kept at a lower temperature (i.e. heating only the head, Fig. 1D). This suggest that the head may not be the most heat sensitive body part (Fig. 5B). When the head was maintained at a lower temperature (abdomen was heated, Fig. 1E), the species differed in response (Fig. 5A). *D. subobscura* and *D. mojavensis* maintained a similar LT_50_ for the abdomen when only the abdomen was heated compared to heating of the whole animal, suggesting that the abdomen is a heat sensitive body part in these two species since selective heating of abdomen gives the same heat tolerance as heating the whole fly. *D. melanogaster* showed a different response as LT_50_ increased in flies when only the abdomen was heated (i.e. a similar response as when the head was selectively heated). This suggest that for *D. melanogaster* both body parts are injured through heat exposure and that the damage may be additive such that it is the total amount of accumulated injury that determines heat tolerance. Overall these experiments showed that the head was not a particular heat sensitive region and the higher LT_50_ values in flies with selective heating of the head suggest that neuronal tissue can survive some degrees beyond the temperature causing SD events.

The increase in LT_50_ for flies with selective heating of the head support the notion that spreading depolarisation is an adaptive mechanism to protect the organism during stress (Robertson, 2004; Rodgers *et al*., 2010). We observed in multiple cases where flies used for the LT_50_ experiments would enter a heat coma (they were completely unresponsive immediately following heat exposure), but they would later resume movement and often recover normal behaviour. Likewise, we observed in the initial behavioural phenotype assays that flies removed from the heat immediately after t/T_coma_ had been observed would recover subsequently. Together these data indicate that SD events are not directly associated with mortality and that nervous failure is not a proximal cause of heat death. Nevertheless, thermal sensitivity of the nervous system could impose a critical challenge to fitness if critical behaviours, such as escape responses, are impaired at stressful temperatures (Montgomery & Macdonald, 1990).

In conclusion, experiments performed for this study show clear interspecific differences in the extent (time/temperature) that the flies can tolerate heat stress, which is related to the overall heat tolerance of the species. Based on the first experiments we find that loss of nervous function is likely to be the cause of the characteristic loss of coordinated movement and coma that is classically used to assess heat tolerance in insects (CT_max_). Our experimental conditions did not allow us to conclude specifically if it is the first or last SD event that is the cause of these phenotypes, and it is also possible that related neuronal failure in other ganglia could play a role. Our second set of experiments with selective heating showed that the head (mainly neuronal tissue) is not particularly heat sensitive compared to other parts of the body. Thus, entry into (reversible) coma and heat mortality are likely different physiological processes and loss of brain function is not the proximal cause of heat death.

The temperature and time span from when the most heat-sensitive species suffered from neural failure to when the CNS of the most heat tolerant species succumbed was large, inviting further studies to investigate adaptations in the CNS to alter heat sensitivity. Our results strongly suggest that hyperthermic loss of CNS function and loss of motor coordination and function (coma) are correlated, which is of clear interest to uncover the physiological perturbations limiting heat tolerance. The role of muscle and neuromuscular synapses in loss of function was not examined in the present study, and although they may also coincide with loss of coordinated movement and heat coma, the correlation between the upstream CNS silencing and loss of function is striking. However, it is also important to appreciate that even small disturbances in nervous function at less stressful temperatures could mean the difference between life and death to an unrestrained animal in nature if its escape response is retarded by nervous dysfunction.

## Acknowledgements

We would like to thank Kirsten Kromand for animal care, Niels U. Kristiansen for help with preparation of thermocouples and Mads K. Andersen for help with the experimental setup and for valuable discussions of the results and experimental design.

## References

Addo-Bediako, A., Chown, S. L. and Gaston, K. J. (2000). Thermal tolerance, climatic variability and latitude. Proc. Biol. Sci. 267, 739–745.

Andersen, M. K. and Overgaard, J. (2019). The central nervous system and muscular system play different roles for chill coma onset and recovery in insects. Comp. Biochem. Physiol. Part A 233, 10–16.

Andersen, J. L., Manenti, T., Sørensen, J. G., MacMillan, H. A., Loeschcke, V. and Overgaard, J. (2015). How to assess *Drosophila* cold tolerance: chill coma temperature and lower lethal temperature are the best predictors of cold distribution limits. Funct. Ecol. 29, 55–65.

Andersen, M. K., Jensen, N. J. S., Robertson, R. M. and Overgaard, J. (2018). Central nervous system shutdown underlies acute cold tolerance in tropical and temperate *Drosophila* species. J. Exp. Biol. 221, jeb179598.

Armstrong, G. A. B., Xiao, C., Krill, J. L., Seroude, L., Dawson-Scully, K. and Robertson, R. M. (2011). Glial Hsp70 protects K+ homeostasis in the *Drosophila* brain during repetitive anoxic depolarization. PLoS One 6, e28994.

Baty, F., Ritz, C., Charles, S., Brutsche, M., Flandrois, J.-P. and Delignette-Muller, M.-L. (2015). A Toolbox for Nonlinear Regression. J. Stat. Softw. 66, 1–21.

Bayley, J. S., Winther, C. B., Andersen, M. K., Grønkjær, C., Nielsen, O. B., Pedersen, T. H. and Overgaard, J. (2018). Cold exposure causes cell death by depolarization-mediated Ca2+ overload in a chill-susceptible insect. Proc. Natl. Acad. Sci. 201813532.

Bigelow, W. D. (1921). The logarithmic nature of thermal death time curves. J. Infect. Dis. 29, 528–536.

Bowler, K. (1963). A Study of the Factors Involved in Acclimatization to Temperature and Death at High Temperatures in *Astacus pallipes* I. Experiments on intact animals. J. Cell. Comp. Physiol. 62, 119–132.

Bowler, K. (1981). Heat death and cellular heat injury. J. Therm. Biol. 6, 171–178.

Bowler, K. (2018). Heat death in poikilotherms: Is there a common cause? J. Therm. Biol. 76, 77–79.

Bowler, K., Duncan, C. J., Gladwell, R.. T. and Davison, T. F. (1973). Cellular heat injury. Comp. Biochem. Physiol. Part A Physiol. 45, 441–450.

Castañeda, L. E., Rezende, E. L. and Santos, M. (2015). Heat tolerance in *Drosophila subobscura* along a latitudinal gradient: Contrasting patterns between plastic and genetic responses. Evolution (N. Y). 69, 2721–2734.

Cossins, A. R. and Bowler, K. (1987). Temperature Biology of Animals. 1st ed. London: Chapman and Hall.

Cowles, R. B. and Bogert, C. M. (1944). A preliminary study of the thermal requirements of desert reptiles. Bull. Am. Museum Nat. Hist. 83, 261–296.

Davison, T. F. and Bowler, K. (1971). Changes in the Functional Efficiency of Flight Muscle Sarcosomes during Heat Death of Adult *Calliphora erythrocephala*. J. Cell. Physiol. 78, 37–48.

Feder, M. E. and Hofmann, G. E. (1999). Heat-shock proteins, molecular chaperones, and the stress response: evolutionary and ecological physiology. Annu. Rev. Physiol. 61, 243–282.

Fox, J. and Weisberg, S. (2011). An R Companion to Applied Regression. Second Edi. Thousand Oaks CA: Sage.

Fraenkel, G. (1960). Lethal High Temperatures for Three Marine Invertebrates: *Limulus polyphemus, Littorina littorea* and *Pagurus longicarpus*. Oikos 11, 171–182.

Friedlander, M. J., Kotchabhakdi, N. and Prosser, C. L. (1976). Effects of Cold and Heat on Behavior and Cerebellar Function in Goldfish. J. Comp. Physiol. A 112, 19–45.

Gaston, K. J. and Chown, S. L. (1999). Elevation and climatic tolerance: a test using dung beetles. Oikos 86, 584–590.

Gladwell, R. T. (1975). Heat death in the crayfish *Austropotamobius pallipes*: Thermal inactivation of muscle membrane-bound ATPases in warm and cold adapted animals. J. Therm. Biol. 1, 95–100.

Gladwell, R. T., Bowler, K. and Duncan, C. J. (1975). Heat death in the crayfish *Austropotamobius pallipes* - Ion movements and their effects on excitable tissues during heat death. J. Therm. Biol. 1, 79–94.

Hamby, R. J. (1975). Heat effects on a marine snail. Biol. Bull. 149, 331–347.

Hazel, J. R. (1995). Thermal Adaptation in Biological Membranes: Is Homeoviscous Adaptation the Explanation? Annu. Rev. Physiol. 57, 19–42.

Heath, J. E. and Wilkin, P. J. (1970). Temperature responses of the desert cicada, *Diceroprocta apache* (Homoptera, Cicadidae). Physiol. Zool. 43, 145–154.

Heath, J. E., Hanegan, J. L., Wilkin, P. J. and Heath, M. S. (1971). Adaptation of the thermal responses of insects. Am. Zool. 11, 147–158.

Jørgensen, L. B., Malte, H. and Overgaard, J. (2019). How to assess *Drosophila* heat tolerance: Unifying static and dynamic tolerance assays to predict heat distribution limits. Funct. Ecol. 33, 629–642.

Jutfelt, F., Roche, D. G., Clark, T. D., Norin, T., Binning, S. A., Speers-Roesch, B., Amcoff, M., Morgan, R., Andreassen, A. H. and Sundin, J. (2019). Brain cooling marginally increases acute upper thermal tolerance in Atlantic cod. J. Exp. Biol. 222, jeb208249.

Kellermann, V., Overgaard, J., Hoffmann, A. A., Flojgaard, C., Svenning, J.-C. and Loeschcke, V. (2012). Upper thermal limits of *Drosophila* are linked to species distributions and strongly constrained phylogenetically. Proc. Natl. Acad. Sci. U. S. A. 109, 16228–16233.

Kimura, M. T. (2004). Cold and heat tolerance of drosophilid flies with reference to their latitudinal distributions. Oecologia 140, 442–449.

Kingsolver, J. G. and Umbanhowar, J. (2018). The analysis and interpretation of critical temperatures. J. Exp. Biol. 221, jeb167858.

Klok, C. J. (2004). Upper thermal tolerance and oxygen limitation in terrestrial arthropods. J. Exp. Biol. 207, 2361–2370.

Koštál, V., Vambera, J. and Bastl, J. (2004). On the nature of pre-freeze mortality in insects: water balance, ion homeostasis and energy charge in the adults of *Pyrrhocoris apterus*. J. Exp. Biol. 207, 1509–1521.

Larsen, E. B. (1943). Problems of heat death and heat injury. Experiments on some species of Diptera. Kgl. Danske Vidensk Selsk. Biol. Medd. 19, 1–52.

Lutterschmidt, W. I. and Hutchison, V. H. (1997a). The critical thermal maximum: data to support the onset of spasms as the definitive end point. Can. J. Zool. 75, 1553–1560.

Lutterschmidt, W. I. and Hutchison, V. H. (1997b). The critical thermal maximum: history and critique. Can. J. Zool. 75, 1561–1574.

MacMillan, H. A. and Hughson, B. N. (2014). A high-throughput method of hemolymph extraction from adult *Drosophila* without anesthesia. J. Insect Physiol. 63, 27–31.

MacMillan, H. A. and Sinclair, B. J. (2011). Mechanisms underlying insect chill-coma. J. Insect Physiol. 57, 12–20.

MacMillan, H. A., Nørgård, M., MacLean, H. J., Overgaard, J. and Williams, C. J. A. (2017). A critical test of *Drosophila* anaesthetics: Isoflurane and sevoflurane are benign alternatives to cold and CO2. J. Insect Physiol. 101, 97–106.

Martinet, B., Lecocq, T., Smet, J. and Rasmont, P. (2015). A Protocol to Assess Insect Resistance to Heat Waves, Applied to Bumblebees (*Bombus* Latreille, 1802). PLoS One 10, e0118591.

Mellanby, K. (1954). Acclimatization and the thermal death point in insects. Nature 173, 582.

Mölich, A. B., Förster, T. D. and Lighton, J. R. B. (2013). Hyperthermic Overdrive: Oxygen Delivery does Not Limit Thermal Tolerance in *Drosophila melanogaster*. J. Insect Sci. 12, 1–7.

Money, T. G. A., Rodgers, C. I., McGregor, S. M. K. and Robertson, R. M. (2009). Loss of Potassium Homeostasis Underlies Hyperthermic Conduction Failure in Control and Preconditioned Locusts. J. Neurophysiol. 102, 285–293.

Montgomery, J. C. and Macdonald, J. a (1990). Effects of temperature on nervous system: implications for behavioral performance. Am. J. Physiol. 259, R191–6.

Neven, L. G. (2000). Physiological responses of insects to heat. Postharvest Biol. Technol. 21, 103–111.

O’Sullivan, J. D. B., MacMillan, H. A. and Overgaard, J. (2017). Heat stress is associated with disruption of ion balance in the migratory locust, *Locusta migratoria*. J. Therm. Biol. 68, 177–185.

Overgaard, J. and MacMillan, H. A. (2017). The Integrative Physiology of Insect Chill Tolerance. Annu. Rev. Physiol. 79, 187–208.

Overgaard, J., Kearney, M. R. and Hoffmann, A. A. (2014). Sensitivity to thermal extremes in Australian *Drosophila* implies similar impacts of climate change on the distribution of widespread and tropical species. Glob. Chang. Biol. 20, 1738–1750.

Pörtner, H.-O. (2001). Climate change and temperature-dependent biogeography: oxygen limitation of thermal tolerance in animals. Naturwissenschaften 88, 137–146.

Prosser, C. L. and Nelson, D. O. (1981). The role of nervous systems in temperature adaptation of poikilotherms. Annu. Rev. Physiol. 43, 281–300.

R Core Team (2018). R: A language and environment for statistical computing.

Robertson, R. M. (2004). Thermal stress and neural function: Adaptive mechanisms in insect model systems. J. Therm. Biol. 29, 351–358.

Robertson, R. M., Dawson-Scully, K. D. and Andrew, R. D. Neural shutdown under stress: an evolutionary perspective. (Submitted).

Robertson, R. M., Spong, K. E. and Srithiphaphirom, P. (2017). Chill coma in the locust, *Locusta migratoria*, is initiated by spreading depolarization in the central nervous system. Sci. Rep. 7, 10297.

Rodgers, C. I., Armstrong, G. A. B., Shoemaker, K. L., LaBrie, J. D., Moyes, C. D. and Robertson, R. M. (2007). Stress preconditioning of spreading depression in the locust CNS. PLoS One 2, e1366.

Rodgers, C. I., Labrie, J. D. and Robertson, R. M. (2009). K+ homeostasis and central pattern generation in the metathoracic ganglion of the locust. J. Insect Physiol. 55, 599–607.

Rodgers, C. I., Armstrong, G. A. B. and Robertson, R. M. (2010). Coma in response to environmental stress in the locust: a model for cortical spreading depression. J. Insect Physiol. 56, 980–990.

Rodríguez, E. C. and Robertson, R. M. (2012). Protective effect of hypothermia on brain potassium homeostasis during repetitive anoxia in Drosophila melanogaster. J. Exp. Biol. 215, 4157–4165.

Sinclair, B. J., Vernon, P., Jaco Klok, C. and Chown, S. L. (2003). Insects at low temperatures: an ecological perspective. Trends Ecol. Evol. 18, 257–262.

Somero, G. N. (1995). Proteins and temperature. Annu. Rev. Physiol. 57, 43–68.

Spong, K. E., Rochon-Terry, G., Money, T. G. A. and Meldrum Robertson, R. (2014). Disruption of the blood-brain barrier exacerbates spreading depression in the locust CNS. J. Insect Physiol. 66, 1–9.

Spong, K. E., Andrew, R. D. and Robertson, R. M. (2016). Mechanisms of spreading depolarization in vertebrate and insect central nervous systems. J. Neurophysiol. 116, 1117–1127.

Stratman, R. and Markow, T. A. (1998). Resistance to thermal stress in desert *Drosophila*. Funct. Ecol. 12, 965–970.

Sunday, J. M., Bates, A. E., Kearney, M. R., Colwell, R. K., Dulvy, N. K., Longino, J. T. and Huey, R. B. (2014). Thermal-safety margins and the necessity of thermoregulatory behavior across latitude and elevation. PNAS 111, 5610–5615.

Terblanche, J. S., Hoffmann, A. A., Mitchell, K. A., Rako, L., le Roux, P. C. and Chown, S. L. (2011). Ecologically relevant measures of tolerance to potentially lethal temperatures. J. Exp. Biol. 214, 3713–3725.

Verberk, W. C. E. P., Overgaard, J., Ern, R., Bayley, M., Wang, T., Boardman, L. and Terblanche, J. S. (2015). Does oxygen limit thermal tolerance in arthropods? A critical review of current evidence. Comp. Biochem. Physiol. A. Mol. Integr. Physiol.

Vorhees, A. S., Gray, E. M. and Bradley, T. J. (2013). Thermal Resistance and Performance Correlate with Climate in Populations of a Widespread Mosquito. Physiol. Biochem. Zool. 86, 73–81.

Wu, J. and Fisher, R. S. (2000). Hyperthermic spreading depressions in the immature rat hippocampal slice. J. Neurophysiol. 84, 1355–1360.

Yi, S. X. and Lee, R. E. (2004). *In vivo* and *in vitro* rapid cold-hardening protects cells from cold-shock injury in the flesh fly. J. Comp. Physiol. B Biochem. Syst. Environ. Physiol. 174, 611–615.

Zachariassen, K. E. (1985). Physiology of cold tolerance in insects. Physiol. Rev. 65, 799–832.

